# Machine learning techniques to derive bioclimatic classifications for Colombia ^☆^

**DOI:** 10.1101/2021.09.05.459033

**Authors:** Richard Rios, Elkin A. Noguera-Urbano, Jairo Espinosa, Jose Manuel Ochoa

## Abstract

Bioclimatic classifications seek to divide a study region into geographic areas with similar bioclimatic characteristics. In this study we proposed two bioclimatic classifications for Colombia using machine learning techniques. We firstly characterized the precipitation space of Colombia using principal component analysis. Based on Lang classification, we then projected all background sites in the precipitation space with their corresponding categories. We sequentially fit logistic regression models to reclassify all background sites in the precipitation space with six redefined Lang categories. New categories were the used to define a new modified Lang and Caldas-Lang classifications.

## 1. Introduction

Bioclimatic classifications seek to divide a study region into geographic areas with similar bioclimatic characteristics (Medina Bermudez and Aldana Buitrago, 2019; IDEAM, 2015). Empiric methods are the most common approaches used to derive bioclimatic classifications by describing relationships between bioclimatic variables, such us temperature or precipitation (Medina Bermudez and Aldana Buitrago, 2019; IDEAM, 2015). Caldas, Lang, and Caldas-Lang classifications are three of the most common classifications derived for Colombia using mean annual values of temperature and precipitation (Medina Bermudez and Aldana Buitrago, 2019; IDEAM, 2015, 2011). For example, Lang classification defines a factor to describe the humidity characteristics of background sites and classify the climate into six categories. This factor is derived as the ratio between the annual precipitation and mean annual temperature. However, this ratio of bioclimatic variables could lead that background sites with low temperatures have similar humidity characteristics than background sites with high level of annual precipitation. As a result, geographical areas like Sierra Nevada de Santa Marta and Chocó could have the same bioclimatic category.

In this document we proposed two bioclimatic classifications for Colombia using machine learning techniques. We initially obtained a modified Lang classification using dimensionality reduction and classification techniques. We then integrated the modified Lang and Caldas classifications to derive a second bioclimatic classification for Colombia.

## 2. Material and methods

### 2.1. Data

The study area encompassed all the continental territory of Colombia. We used five bioclimatic variables at 1 − *km*^2^ spatial resolution, which were: annual mean temperature (*Bio*_1_), annual precipitation (*Bio*_12_), precipitation of wettest month (*Bio*_13_), precipitation of driest month (*Bio*_14_), precipitation seasonality (*Bio*_15_)(Fick and Hijmans, 2017).

### 2.2. Caldas Classification

The Caldas classification relates the variability of the mean annual temperature with the altitude, making it only valid for regions located in the tropical zones (IDEAM, 2015). Figure B.4 of supplemental material depicts the Caldas classification for Colombia. Bioclimatic categories are provided in Table 2.

**Table 1:**
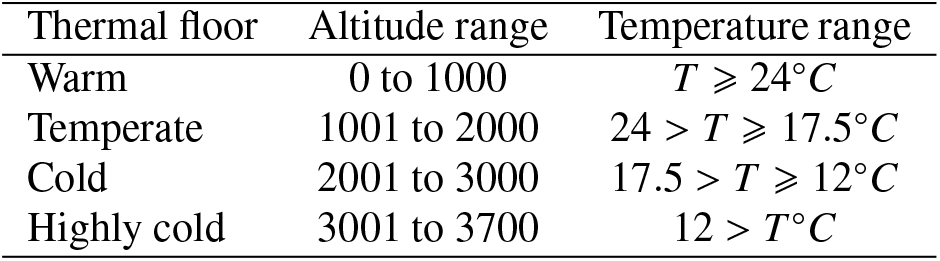
Caldas classification. Abbreviations: T, temperature

| Thermal floor | Altitude range | Temperature range |
| --- | --- | --- |
| Warm | 0 to 1000 | $T \geq 24^\circ C$ |
| Temperate | 1001 to 2000 | $24 > T \geq 17.5^\circ C$ |
| Cold | 2001 to 3000 | $17.5 > T \geq 12^\circ C$ |
| Highly cold | 3001 to 3700 | $12 > T^\circ C$ |

**Table 2:**
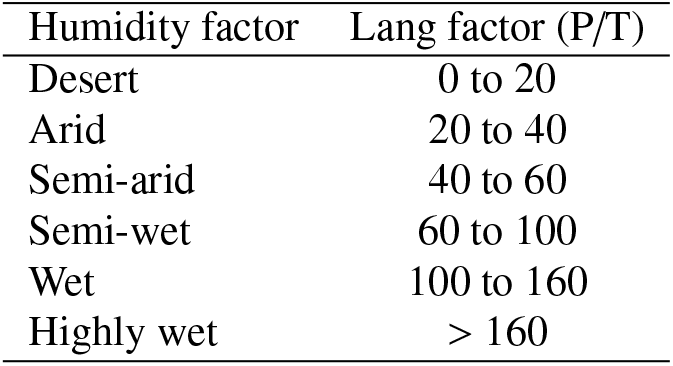
Lang classification. Abbreviations: P, precipitation; T, temperature

### 2.3. Lang Classification

Lang classification uses the ratio between the annual precipitation [*mm*] and mean annual temperature [°*C*] to describe the humidity characteristics of background sites (IDEAM, 2015). This ratio called Lang factor is then used to classify the climate into six categories, see Table 3. Figure B.5 of supplemental material depicts the Lang classification for Colombia.

**Table 3:**
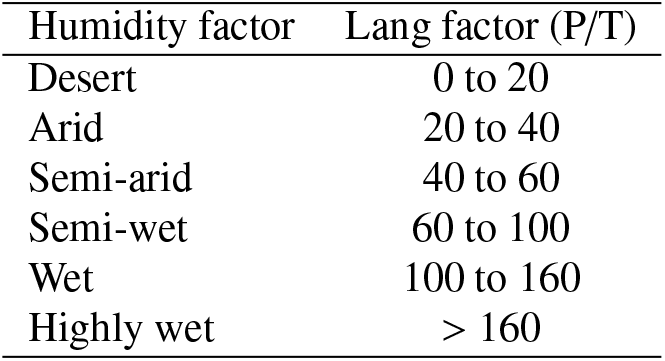
Lang classification. Abbreviations: P, precipitation; T, temperature

### 2.4. Modified Lang Classification

We firstly obtained a characterization of the Colombia’s precipitation space by applying a principal component analysis (PCA) to the set of precipitation variables, using standard procedures (Abdi and Williams, 2010). We then calculated the Lang factor for each background site and determined its corresponding bioclimatic category.

Later, we projected all background sites with their bioclimatic categories into the precipitation space, and based on these categories, we sequentially fit logistic regression models to redefine the Lang classification’s blioclimatic categories (Bishop, 2006; Hastie et al., 2017). We assumed that two adjacent or closer background sites in the precipitation space should share similar bioclimatic characteristics.

#### Characterization of precipitation space

For the set of precipitation variables, the first two principal components (PC)s contributed 96.8% of the variability. Fig 1 depicts the loadings of the precipitation variables projected in the latent space spanned by the first two PCs, hereby name precipitation space. It can be observed, for instance, that negative and positive values of the first PC are related to areas with high and low levels of precipitation, respectively. Meanwhile, positive values of the second PC are related to areas with high variability of monthly precipitation over the course of the year. Fig 2 depicts the projection of all background sites in the precipitation space with their corresponding bioclimatic categories. Then, it can be observed that background sites classified as highly wet (blue dot points) were distributed along the first PC: some highly wet sites were even close to dry and desert categories, see green and red colors in Figure 2, respectively.

**Figure 1:**
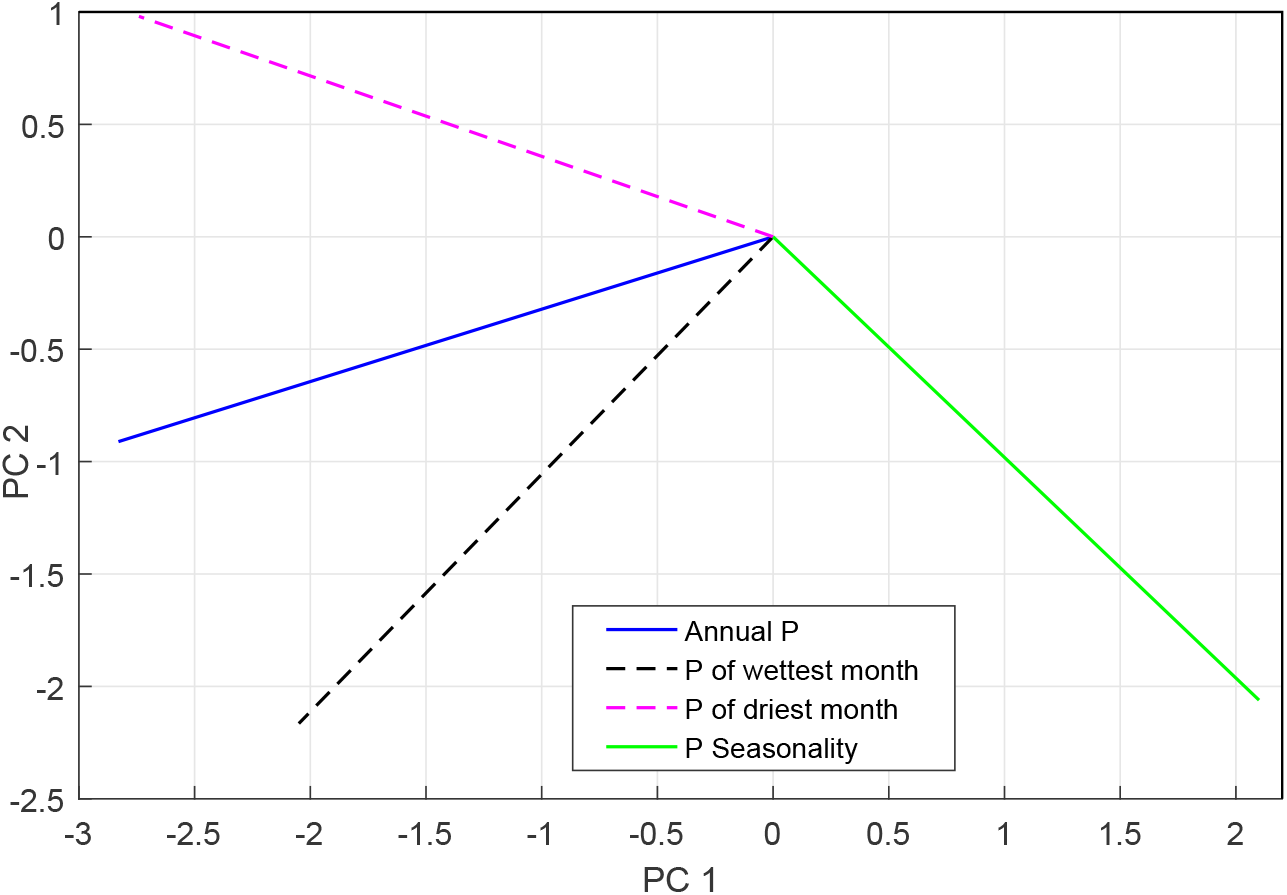
Projection of the precipitation variables in the first two PCs. Abbreviations: P, precipitation.

**Figure 2:**
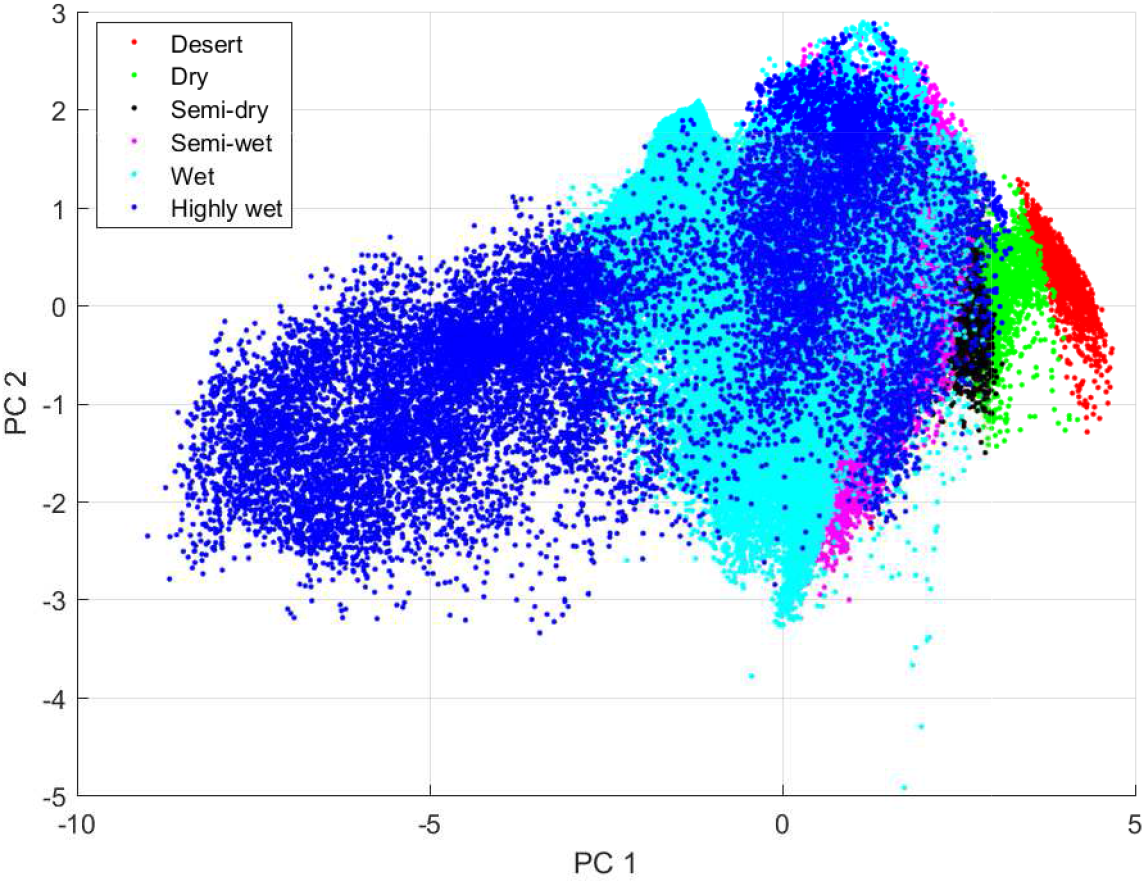
Projection of all background sites in the first two PCs with their corresponding categories in the Lang classification. This projection represent the characterization of the Colombia’s precipitation space, whose axes are given by the first two PCs. Abbreviations: P, precipitation.

#### Classification of precipitation space

We sequentially fit a set of classifiers to divide the precipitation space in six categories. We used the Lang classification’s categories as labels to define a multi-class classification problem. All classifiers were fit by using regularized logistic regression, see details in the supplement material. We described as follows the step performed to derive a modified Lang classification :

1. Based on the Lang classification, we fit a logistic model to classify dry (class 0) and desert (class 1) background sites. Then, this model was used to classify all background sites between 0 or 1. We redefined the bioclimatic category of desert with all background sites assigned to the class 1.
2. Based on the semi-dry, dry, and redefined desert categories, we fit a logistic model to separate semi-dry background sites (class 0) from dry and desert background sites (class 2). Then, this model was used to classify all background sites between 0 or 2. We redefined the bioclimatic category of dry with all background sites assigned to the class 2 and class 1 (from step 1).
3. We then fit two logistic regression models to separate semi-wet background sites (class 0) from semi-dry, dry, and desert background sites (class 3). Then, these models were used to assign all background sites to class 0 or 3. We redefined the bioclimatic category of semi-dry with all background sites assigned to the class 3 and class 2 (from step 2).
4. Similarly, we fit two logistic regression models to separate wet background sites (class 0) from semi-wet, semi-dry, dry, and desert background sites (class 4). Then, these models were used to assign all background sites to class 0 or 4. We redefined the bioclimatic category of semi-wet with all background sites assigned to the class 4 and class 3 (from step 3).
5. Finally, we fit a logistic model to separate highly wet background sites (class 0) from wet and semi-wet background sites (class 5). Then, this model was used to assign all background sites to class 0 or 5. We redefined the bioclimatic category of wet with all background sites assigned to the class 5 and class 4 (from step 4). Similarly, we redefined the bioclimatic category of highly-wet with all background sites assigned to the class 0.

Parameters tuned for the classifiers are provided in Table S1. Meanwhile, Figure 3 depicts the projections of all background sites in the precipitation space with their corresponding new redefined bioclimatic category. Similarly, Figure B.6 in the supplemental material depicts the modified Lang classification for Colombia by re-projecting the background sites to the geographical space. As a result, it can be observed that highly-wet background sites were mostly located in the Pacific region of Colombia, which has high-level of annual precipitations. Meanwhile, for instance, areas like Sierra Nevada de Santa Marta or Parque Nacional Natural el Cocuy were recategorized as semi-wet, semi-arid, and arid.

**Figure 3:**
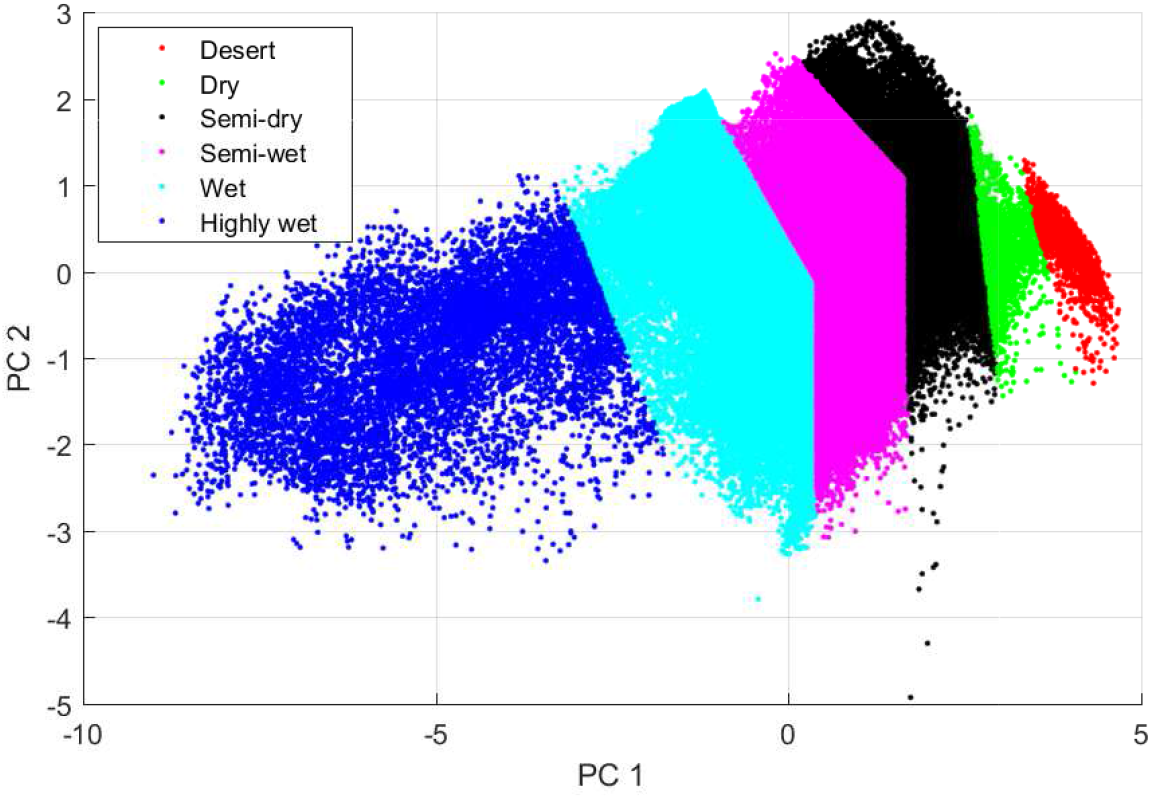
Projection of all background sites in the first two PCs with their corresponding redefined categories. Logistic regression models were used to classify all background sites into the six categories. Abbreviations: P, precipitation.

### 2.5. Modified Caldas-Lang Classification

Temperature and humidity factor are two important covariates used to describe the climate in tropical regions (IDEAM, 2015; Medina Bermudez and Aldana Buitrago, 2019). Caldas-Lang classification is derived by combining both Caldas and Lang Classification. Categories are defined by taking a first label from the Caldas classification and a second label from the Lang Classification. We followed the same procedure to derive a modified Lang-Caldas classification by integrating the Caldas and modified Lang classification. Figure B.7 in the supplemental material depicts the modified Caldas-Lang classification for Colombia.

## 4. Acknowledgements

This research was supported by the Departamento Administrativo de Ciencia, Tecnología e Innovación (COLCIENCIAS) – Colombia with the “Programa de Estancias Postdoctorales Beneficiarios COLCIENCIAS 2017” grant.

## Appendix A.

### Regularized logistic regression model

Considering a set of *n* records represented in a *M*−dimensional feature space as *w*_*i*_(*i* = 1, …, *n*), the goal was to determine a logistic regression model that optimally separate records between class 1 and 0. This model is described as follows:

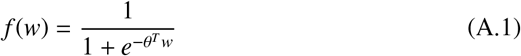

where *θ* = [*θ*_1_, …, *θ*_*m*_]^*T*^ was the vector of model parameters, and *w =* [*w*_0_, …, *w*_*m*_]^*T*^ was the vector of features, holding that *w*_0_ = 1. The function *f* then predicted the probability that *y* = 1 on the input vector *w*, i.e., *f* (*w*) = *P*(*y* = 1*/θ, w*). As a result, given a feature vector *w*, the function *f* assigned it to the class 1 if *θ*^*T*^ *w* ⩾ 0.5 while to class 0 otherwise.

For a training set (*w*^(*i*)^, *y*^(*i*))^, where *i* = 1, …, *n*, we sought to find the vector of parameters *θ* and *λ* that minimize the following cost function:

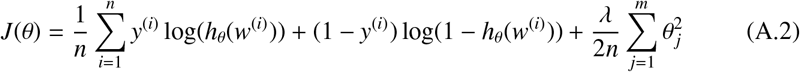

where *λ* is the regularization parameter. The following table provides the parameters tuned for the logistic regression models.

**Table A.4:**
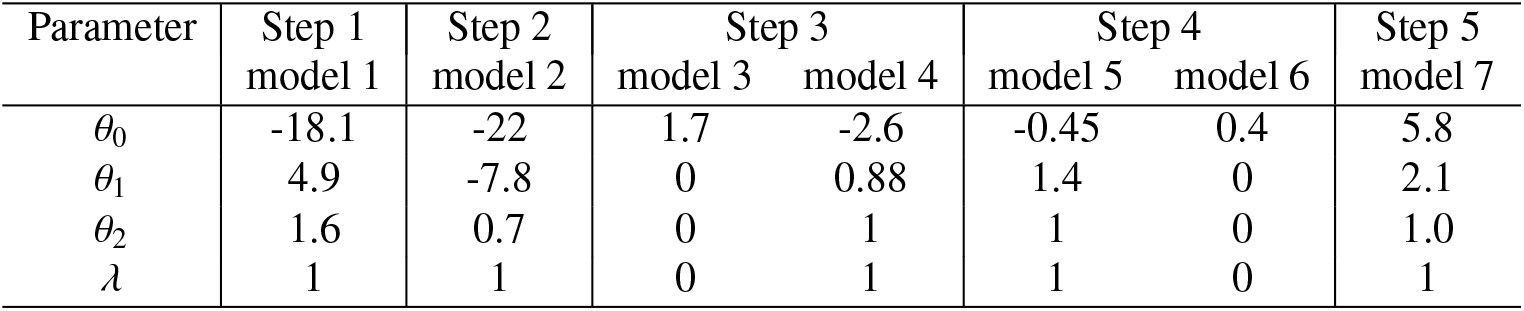
Parameters of the logistic regression models used to derive the modified Lang classification.

## Appendix B.

### Bioclimatic classifications

All bioclimatic classification are available at the Zenodo repository https://doi.org/10.5281/zenodo.5396516

**Figure B.4:**
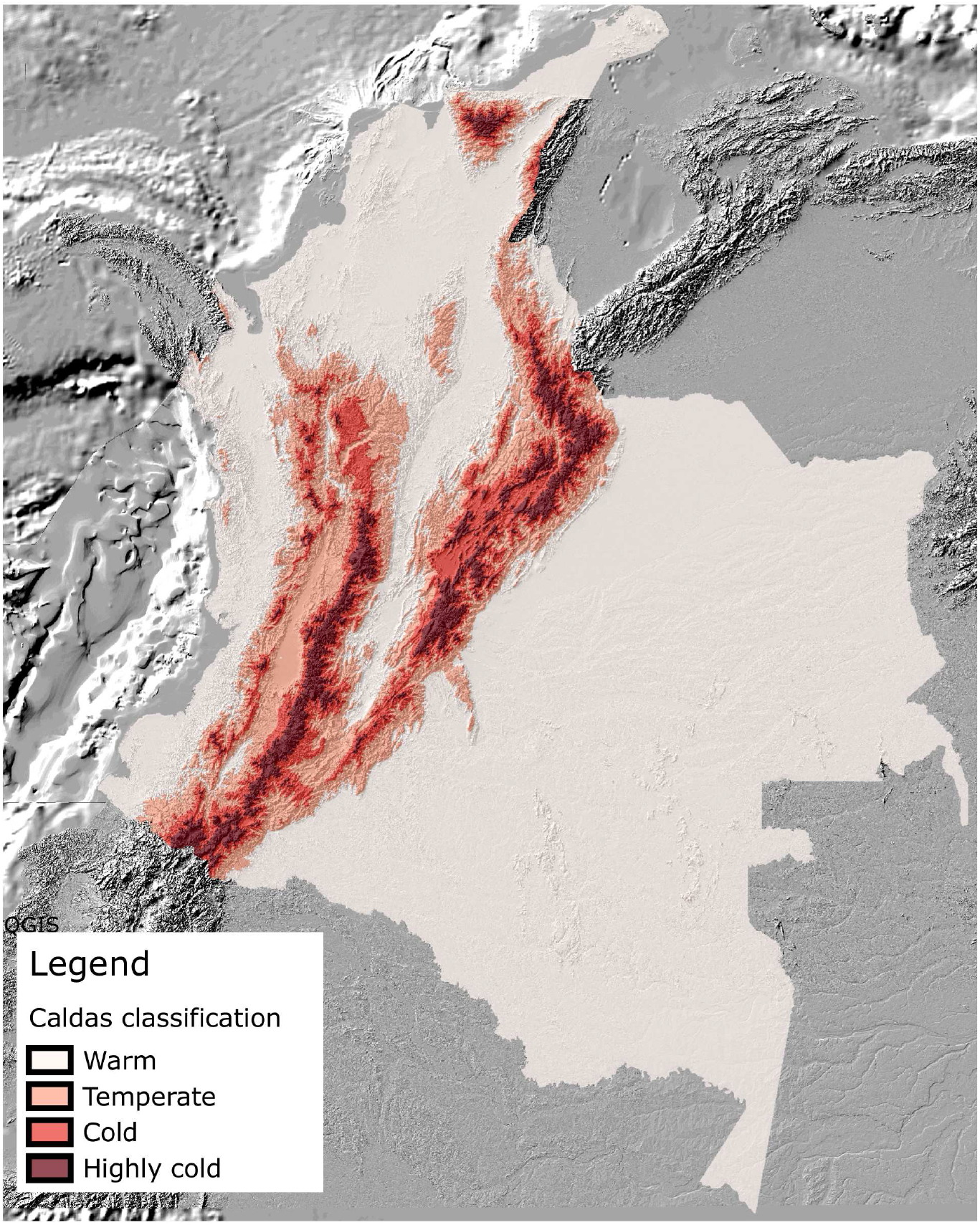
Caldas classification

**Figure B.5:**
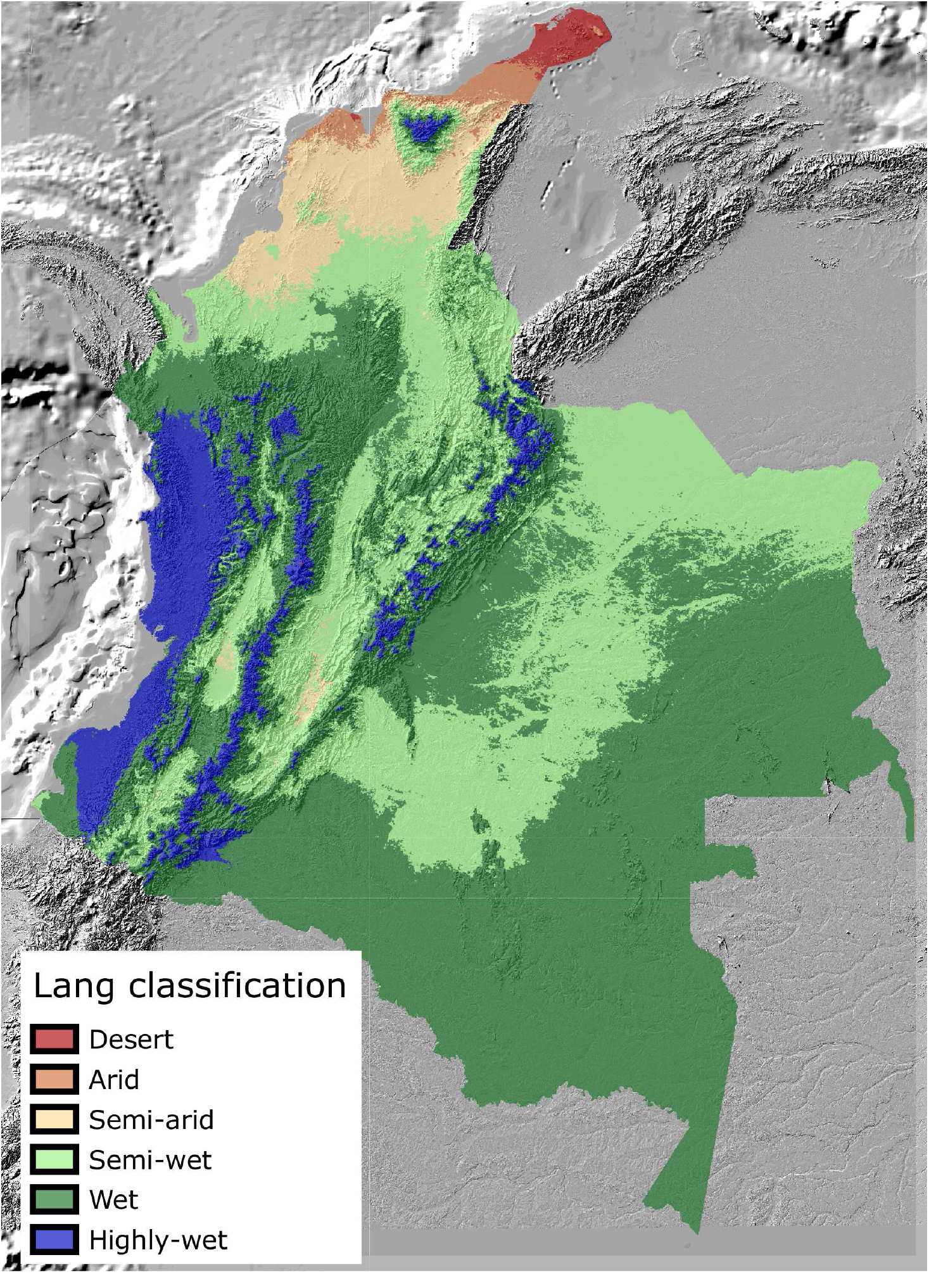
Lang classification

**Figure B.6:**
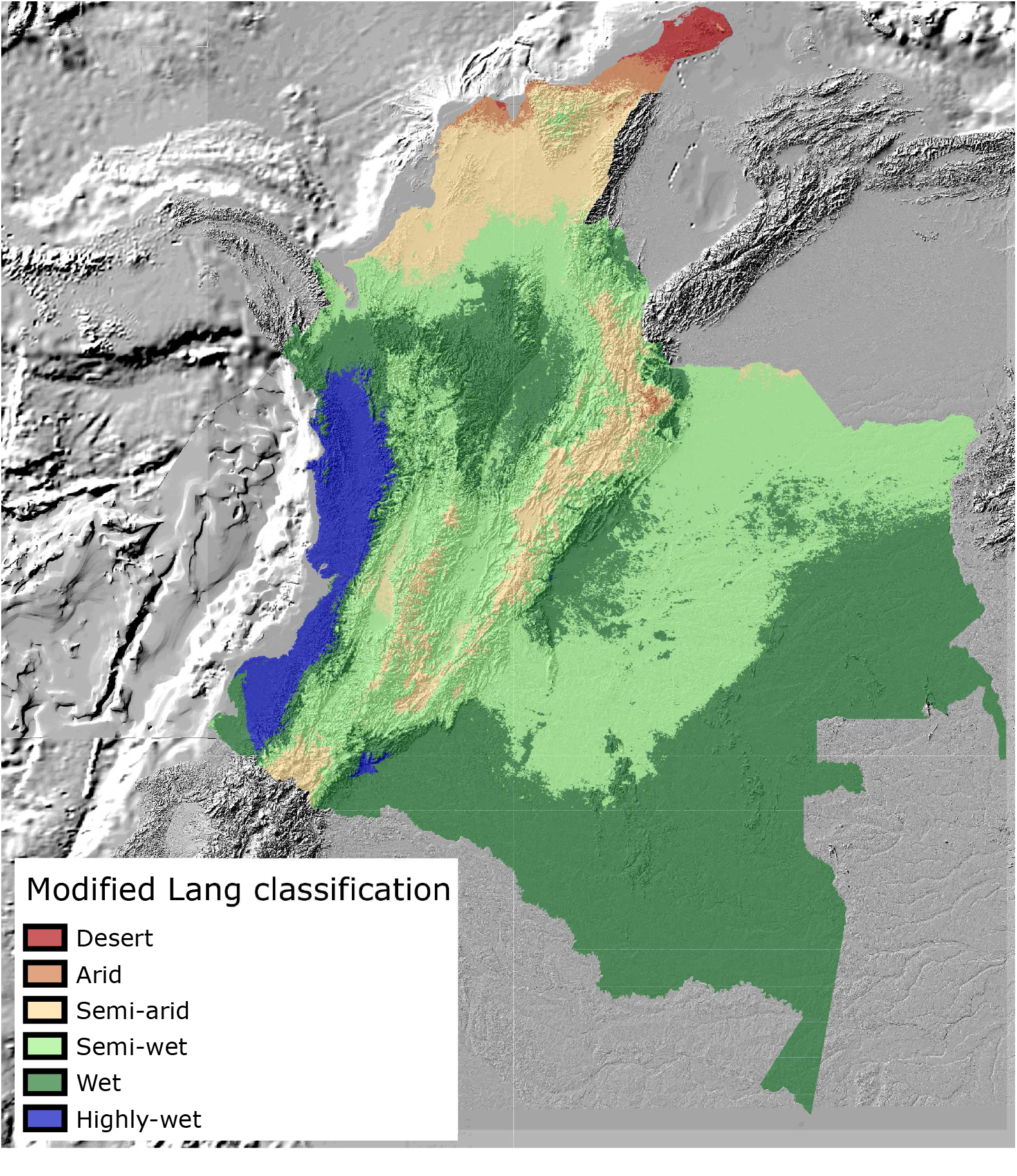
Modified Lang classification

**Figure B.7:**
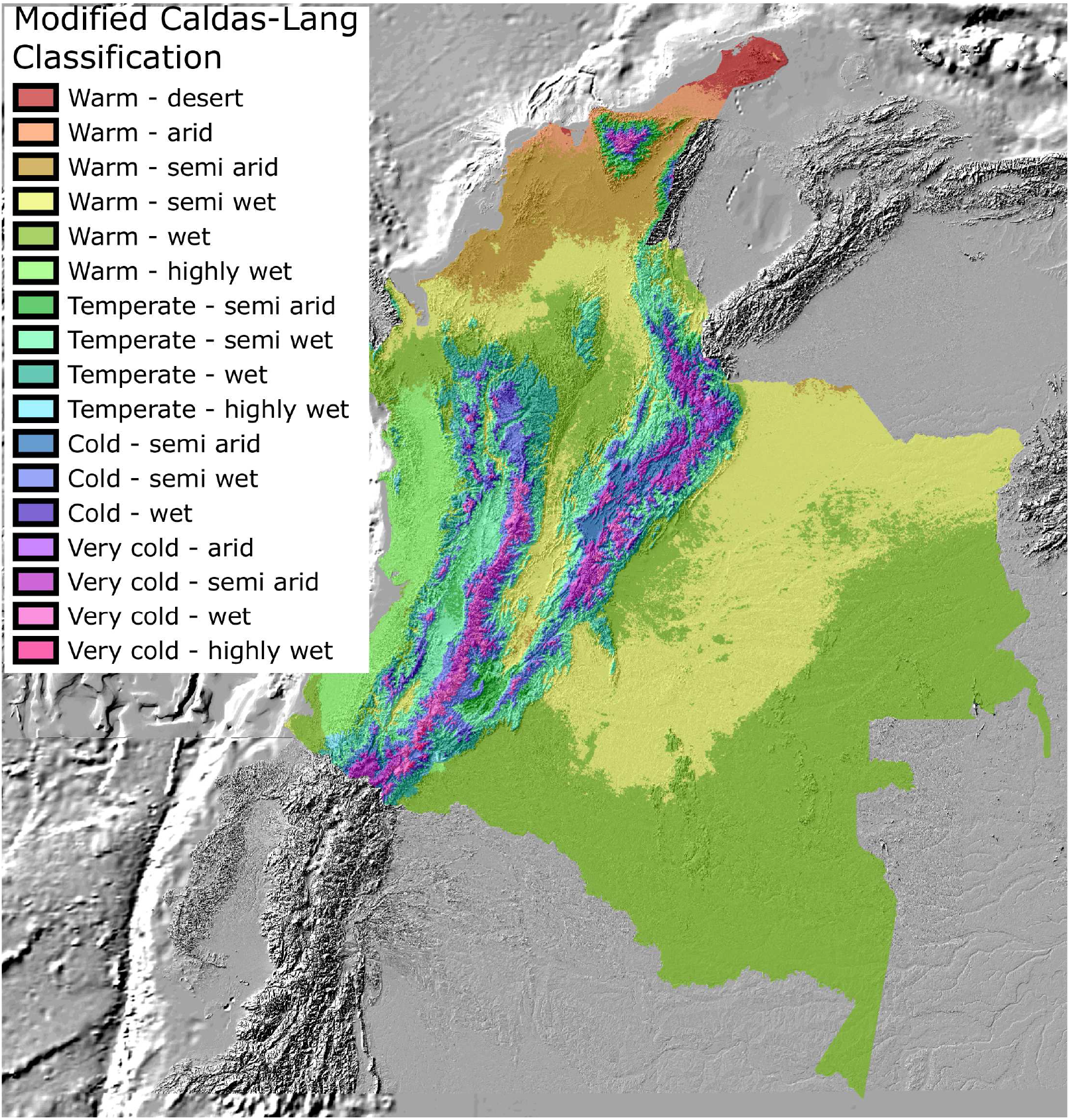
Modified Caldas-Lang classification

